# SMYD1-mediated Mono-Methylation of Lysine K35 of the sarcomeric Myosin Heavy Chain (MHC) is fundamental for thick filament assembly in zebrafish and human iPSC-derived cardiomyocytes

**DOI:** 10.1101/2024.03.27.585692

**Authors:** Federica Diofano, Chidinma Amadi, Bernd Gahr, Karolina Weinmann, Wolfgang Rottbauer, Steffen Just

## Abstract

The SMYD family is a unique class of lysine methyltransferases (KMTases) known to methylate histones but also non-histone proteins. Among the five SMYD family members (1-5), SMYD1 was identified as a heart- and skeletal muscle-specific KMTase, which, together with Unc45b and Hsp90a, interacts with Myosin thereby regulating thick filament assembly. However, the process by which SMYD1 orchestrates Myosin assembly is largely unknown. Here, we found that SMYD1 physically interacts with Myosin heavy chain (Myh) at its N-terminus and that the Myh N-terminus specifically gets mono-methylated by SMYD1 at lysine 35 (K35). Accordingly, methylated Myh is properly integrated into functional sarcomeres, whereas unmethylated Myh molecules in Smyd1-deficient zebrafish are efficiently degraded by the Ubiquitin Proteasome System (UPS) leading to defective thick filament assembly. Although the inhibition of the UPS by MG132 is able to reconstitute Myosin levels in Smyd1-deficient zebrafish embryos, thick filament assembly is still blocked due to the lack of K35 Myh mono-methylation. Similar to the situation in zebrafish striated muscle cells, SMYD1-mediated MYH methylation is also critical for thick filament assembly in human cardiomyocytes, indicating cross-species conservation of this fundamental mechanism of Myosin methylation, which has been first described about 40 years ago. Further investigations will now be essential to explore the therapeutic potential of targeting this pathway in cardiomyopathies and skeletal muscle disorders.

## Introduction

Proper contractile function of both, cardiomyocytes and skeletal muscle cells relies on the coordinated synthesis, renewal, and assembly of numerous contractile, structural, and regulatory proteins into sarcomeric units, a process known as sarcomerogenesis. Despite its importance, much remains to be unravelled about the regulatory mechanisms of sarcomere assembly during development and its fine-tuning during sarcomere renewal.

SMYD proteins that contain SET [SU(VAR)3-9, Enhancer of zeste and Trithorax] and MYND (Myleoid, Nervy and DEAF-1) domains are predominantly involved in the regulation of gene transcription by the methylation of lysine residues on histone tails (histone methyltransferase activity) and thereby the remodelling of chromatin [1–3]. In addition to their role in histone modification, SMYD proteins can also regulate functions of non-histone proteins through site-specific lysine methylation of target proteins (protein lysine methyltransferase)[4, 5]. For instance, in striated muscle cells, Hsp90 is specifically methylated by the methyltransferase Smyd2 which enables the formation of a protein complex containing Smyd2, Hsp90, and the elastic filament Titin [6]. The absence of Smyd2 in turn leads to a loss of Hsp90 methylation, compromised Titin stability and impaired muscle function [6]. Additionally, the striated muscle specific methyltransferase Smyd1 was shown to physically interact with Myosin and thereby to play an essential role in Myosin thick filament assembly [7]. The modification of Myosin by methylation has been known for more than 40 years, but the functional role of this modification and the proteins that transfer methyl groups to Myosin are still unknown [8]. Remarkably, in hypertrophic cardiomyopathy (HCM) affecting 1 in 500 people, about 40 percent of HCM-causing mutations are found in the human cardiac Myosin gene. Whether aberrant methylation of Myosin is also involved in HCM development and/or progression is completely unknown. In this study, we demonstrate that the methyltransferase Smyd1 predominantly localized to the cytosol and physically interacts with sarcomeric Myosin heavy chain (Myh) at its N-terminus. We identified a specific N-terminal site for Myh lysine methylation (K35) and proved that lysine K35 methylation is critically dependent on Smyd1 function. In zebrafish, we found that methylated Myh is effectively integrated into functional sarcomeres whereas unmethylated Myh is destabilized and degraded by the Ubiquitin Proteasome System (UPS). Similar to zebrafish striated muscle cells, Adeno-associated virus (AAV)-mediated knock-down of SMYD1 in human iPSC-derived cardiomyocytes (hiPSCMs) led to the loss of K35 Myh methylation, Myh degradation and ultimately defective sarcomerogenesis, demonstrating the evolutionary conservation of Smyd1-mediated Myh methylation and its critical role in thick filament assembly. In summary, our data demonstrates for the first time that Myh methylation at lysine 35 is specifically processed by Smyd1 and decodes the 40-year-old mystery of the *in vivo* role of Myosin methylation in striated muscle cells.

## Results

### Sarcomeric Myosin Heavy Chain is mono-methylated at N-terminal Lysine 35

The methyltransferase Smyd1b was shown to physically interact with Myosin in zebrafish striated muscle cells [7, 9]. Although it was known for decades that Myosin heavy chain (Myh) proteins are methylated, neither the methyltransferase mastering the methylation of specific, so far undescribed, Myh lysine (K) residues nor the biological regulatory function of this post-translational modification was known. Hence, to assess whether Myh-interacting methyltransferase Smyd1 is involved in the site-specific methylation of Myh, we first conducted a Mass Spectrometry (MS)-based proteomic analysis of the protein methylation status in Smyd1b-deficient *flatline* (*fla)* mutant zebrafish embryos compared to wild-type clutchmates at 72 hours post fertilization (hpf). The analysis of the methyl-K sites on the peptides corresponding to the proteins recognized in the MS analysis identified around 100 proteins with differentially regulated methylation sites (log fold change >2 and <-2) (Suppl. Table 1). Interestingly, among the proteins with most significantly downregulated methyl-K sites, we found different isoforms of sarcomeric Myh (Fig. 1A), strongly suggesting that Smyd1b is the methyltransferase responsible for the posttranslational modification of Myh via lysine methylation. Additionally, our MS-analysis clearly demonstrated the mono-methylation of Myh peptides at the N-terminal lysine 35 (K35). The methylation of Myh at residue K35 was confirmed by analysing the fragmentation spectra of the Myh peptide ERVEAQUNKPFDAK corresponding to amino acids 23 to 35, which was found methylated in wild-type clutchmates and non-methylated in Smyd1b-deficient *fla* mutants (Fig. S1 and Suppl. Table 1). As shown in Supplemental Figure 1, we were able to confirm mono-methylation on K35 as we detected b8-, b11-, y2- and y5-ions in the respective MS2-spectrum mapping the methylation to K35 and not K30 (Fig. S1). Notably, the N-terminal region/sequence of the muscle-specific Myh is highly conserved across various species, including human, mouse, rabbit, *Danio rerio, Drosophila,* and *C.elegans* (Fig. 1B), implying the evolutionary conservation of this posttranslational modification across vertebrate and invertebrate species. The N-terminus plays a crucial role in mediating the conformational changes that occurs within the Myh head during Actin binding [10, 11].

**Figure 1.**
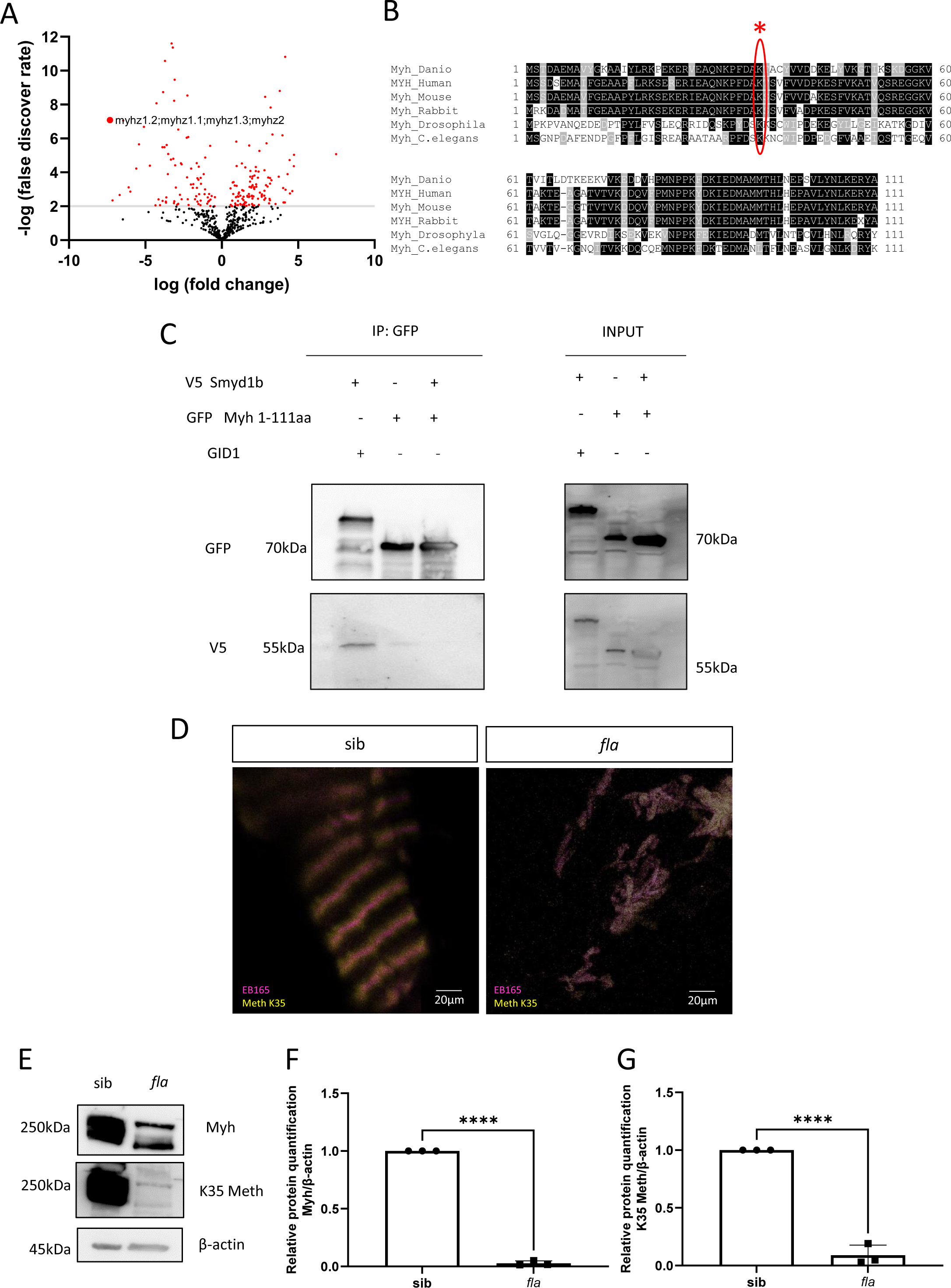
Loss of Smyd1 leads to reduced methylation of its interactor MYH at lysine 35. **(A)** Volcano Plot of MS data showing the methylation profile in the proteome of Smyd1b-deficient *fla* mutant embryos. Methylation of sarcomeric Myosin Heavy Chains (Myh) are significantly reduced in *fla* mutants. (**B**) Myh alignment between species showing that the N-terminus of Myh is highly conserved between species. Methylated lysine residue K35 is highlighted by an asterisk. (**C**) Immunoprecipitation assay using V5-tagged Smyd1b and GFP-tagged N-terminal Myh (1-111aa), demonstrating Smyd1b and Myh interaction. GID1 was used as control. (N=4) (**D**) Immunostaining of Smyd1b-deficient *fla* mutants at 72hpf, showing loss of Myh (magenta) accompanied by the loss of sarcomeric structures and the reduction of methylated K35 Myh (yellow). **(E)** Immunoblot analysis of Smyd1b-deficient *fla* mutant protein lysate compared to protein lysate obtained from wild-type clutchmates (indicated as sib) at 72 hpf (N=3). Total Myh and K35 Myh methylation are reduced in *fla* mutants **(F)** Quantification demonstrates significantly reduced Myh protein levels in *fla* mutants compared to wild-type clutchmates (N = 3; mean ± S.D, P ≤ 0.0001 determined using two-tailed t-test. Error bars indicate s.d.; **** P < 0.0001). ß-Actin was used as a loading control. **(G)** Methylated lysine 35 (K35 Meth) Myh protein levels are significantly reduced in *fla* mutants compared to clutchmates (N = 3; mean ± S.D, P ≤ 0.0001 determined using two-tailed t-test. Error bars indicate s.d.; **** P < 0.0001). ß-Actin was used as a loading control.

Next, to prove that Smyd1b physically bind to the N-terminus of Myh, we performed an immunoprecipitation assay using V5-tagged Smyd1b and GFP-tagged N-terminal (1-111aa) Myh. After overexpression of both constructs in HEK 293 cells we were able to co-immunoprecipitate Smyd1b and N-terminal Myh thereby confirming their physical interaction further substantiating the role of Smyd1 in the methylation of Myh (Fig. 1C).

To be able to assess the biological role of K35 Myh methylation in more detail, we generated a polyclonal Myh antibody that selectively recognizes the monomethylated K35 residue. By investigating the cytosol-localization of K35-methylated Myh in wild-type but also Smyd1b-deficient *fla* zebrafish embryos, we found that K35-methyl-Myh localized exclusively at the sarcomere, suggesting that K35-methylation is critical for proper thick filament assembly into functional sarcomeric units. By contrast, K35-methyl-Myh was undetectable in Smyd1b-deficient *fla* zebrafish embryos as shown by immunostainings and Western Blot analyses (N=3, P<0.0001) (Fig. 1D, E, F, G). These findings imply that Smyd1b methylates Myosin *in vivo* and that K35-specific Myosin mono-methylation is necessary for sarcomeric thick filament assembly *in vivo*.

### The Methyltransferase SMYD1 specifically methylates MYH at Lysine 35

Finally, to biochemically prove that SMYD1 is the protein methyltransferase that is methylating Myosin at lysine 35 and that this biological phenomenon is conserved between zebrafish and humans, we performed a methyltransferase assay utilizing human SMYD1 and MYH and subsequently assessed K35-MYH methylation by using our custom-designed K35-methyl-Myh antibody. We precipitated SMYD1 and N-terminal MYH that had been incubated in methylation buffer in presence and absence of the methionine donor (S-5’-Adenosyl)-L-methionine) (Fig. 2A). As anticipated, the methylated K35 antibody signal was only detected in the presence of donor methionine with MYH, whereas no signal was detected in the absence of methionine (Fig. 2A). Demonstrating that the methylation of human MYH is dependent on the function of SMYD1.

**Figure 2.**
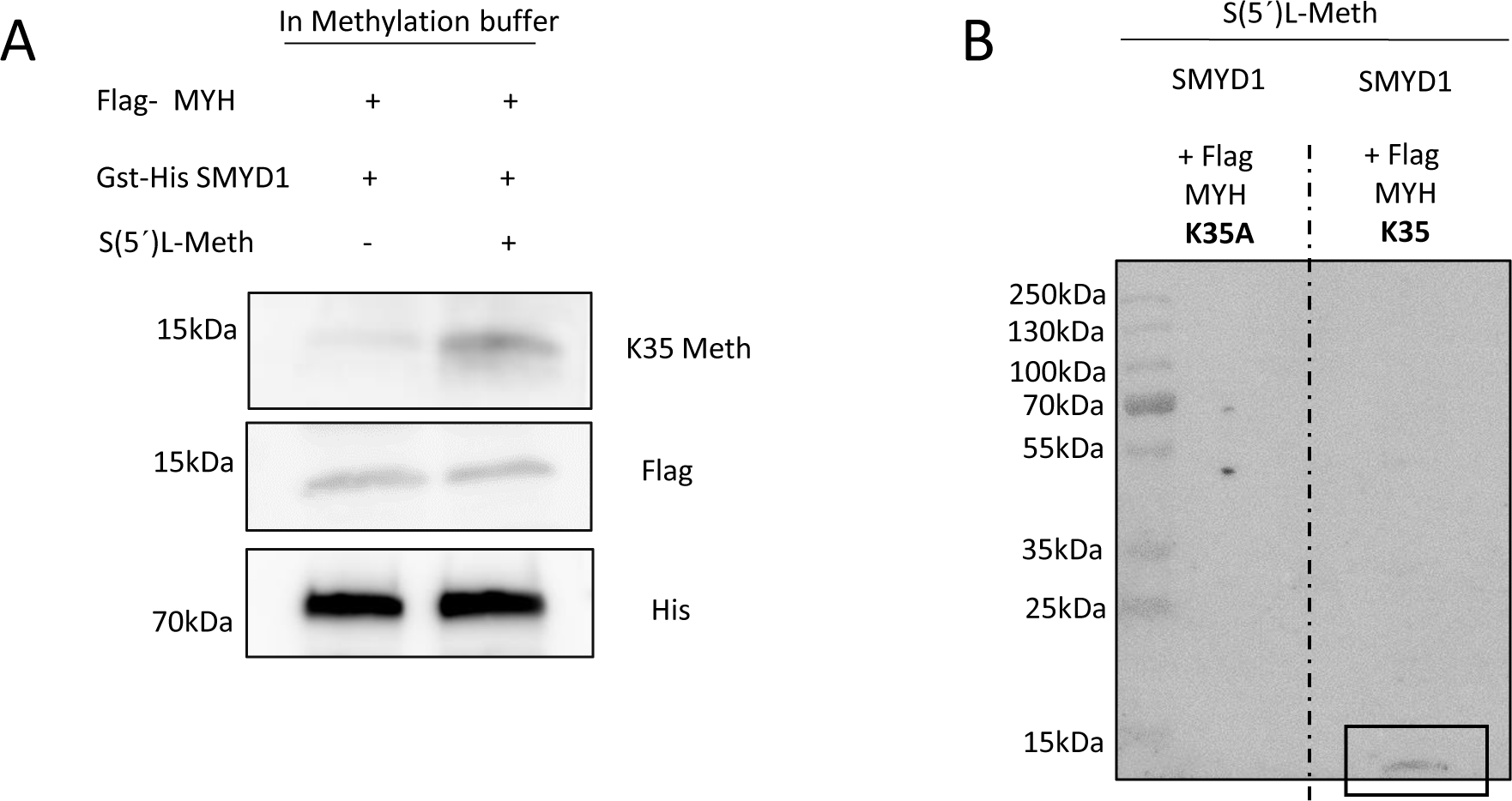
SMYD1 methylates MYH at lysine residue 35. **(A)** *In vitro* methyltransferase assay using GST-His SMYD1 and Flag-MYH (1-111aa), shows that MYH K35 is methylated by SMYD1 in the presence of S-5’Adenosyl-L-methionine (S(5’)L-Meth) (N=3). **(B)** *In vitro* methyltransferase assay using wild-type N-term MYH (1-111aa) and methylation-resistant-N-term MYH K35A (lysine to alanine at position 35). Methylation was only detectable in wild-type N-term MYH but not in the Myosin K35A peptide (N=3).

Furthermore, to demonstrate the ability of SMYD1 to mono-methylate MYH at the Lysine residue K35, we performed an immunoprecipitation assay using GST-His-SMYD1 and the two Flag-tagged N-terminal MYH1 variants, unmodified (wild-type) and methylation-deficient K35A (Lysine35Alanine) (Fig. 2B and S2). Remarkably, the K35 antibody signal was only detected with the fragment containing methylation-accessible K35, confirming that SMYD1 specifically methylates the K35 residue of MYH1 (Fig. 2B).

This biochemical assay proves that SMYD1 indeed mono-methylates the K35 residue in MYH1.

### Myh Methylation is critical for Thick Filament Assembly and loss of Myh Methylation results in its effective Proteasomal Degradation

Next, we aimed on the definition of the biological relevance of Smyd1-mediated Myh K35 methylation *in vivo*. As recently shown by us and others, Smyd1 deficiency leads to the massive degradation of misfolded Myosin proteins, most likely by the ubiquitin proteasome system (UPS) [7, 9, 12]. Whether Myh K35 methylation is critical for proper Myosin folding or involved in the protection of Myosin from UPS-mediated degradation is unknown. In this context, it is interesting to note that methylated lysine residues can regulate protein turnover by preventing ubiquitination [13].

To test the hypothesis that unmethylated Myosin is efficiently ubiquitinylated and subsequently degraded thereby leading to defective thick filament assembly, we first measured the level of protein ubiquitination in Smyd1-deficient *fla* zebrafish embryos by Western blotting. Interestingly, the levels of ubiquitinylated proteins were significantly increased in Smyd1-deficient zebrafish (N=3, P=0.0080) (Fig. 3A, B), implying that more proteins, including Myosin, are prone for degradation by the UPS. To investigate whether UPS function is compromised by the increase of degradation-prone proteins, we next measured the levels of ubiquitinylated proteins in *fla* mutants and sibling after treatment with the established UPS inhibitor MG132. We found that MG132 treatment led to a continued increase of ubiquitinylated proteins in *fla* mutants and wild-type, excluding compromised UPS function as the cause of defective thick filament assembly (N=3, P=0.9043) (Fig. 3C, D).

**Figure 3.**
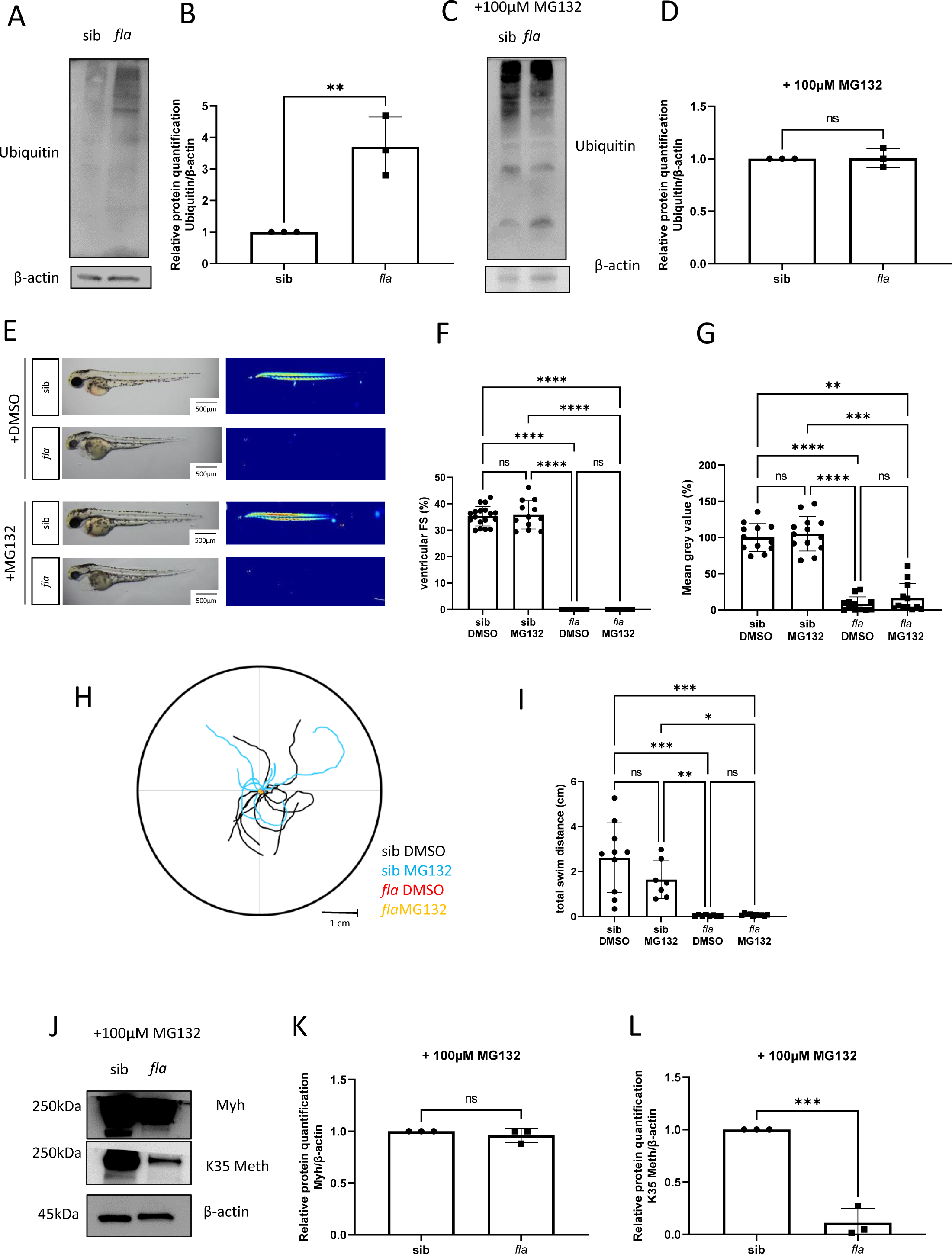
Defective K35 Myh methylation leads to its proteosomal degradation. **(A, B)** *fla* embryos show an accumulation of ubiquitinylated protein compared to wild-type siblings (N=3). Ubiquitin protein quantification shows an increase of ubiquitinylated proteins in *fla* mutants compared to wild-type siblings (N = 3; mean ± S.D, P =0.0080 determined using two-tailed t-test. Error bars indicate s.d.; ** P < 0.01). ß-Actin was used as a loading control. **(C)** UPS inhibition by MG132 treatment leads to a continued increase of ubiquitinylated proteins in *fla* mutants (compared to untreated *fla* mutants **(A)**), demonstrating that UPS function in not compromised in Smyd1-deficient *fla* mutants (N=3). **(D)** Quantification of ubiquitinylated protein, after MG132 treatment, shows no differences in ubiquitinylated proteins in *fla* mutants compared to wild-type siblings (N = 3; mean ± S.D, P=0.9043 determined using two-tailed t-test. Error bars indicate s.d.; ns P >0.05). ß-Actin was used as a loading control. **(E)** Lateral view of brightfield and birefringence images for wild-type and *fla* mutant embryos at 72 hpf, DMSO and MG132 treated, respectively. **(F)** Quantification of ventricular FS at 72 hpf reveals absence of contraction in DMSO- and MG132-treated *fla* embryos (N = 3, n = 4/5, mean ± S.D. One-way ANOVA followed by Kruskal-Wallis multiple comparison analysis. Error bars indicate s.d.; **** P < 0.0001; ns, not significant). **(G)** Densitometric analysis of the birefringence signal at 72 hpf reveals significantly reduced signal intensity in both, DMSO- and MG132-treated *fla* mutant embryos, implying the disorganization of skeletal muscle (N = 3, n = 4/5; P = 0.0001; One-way ANOVA followed by Kruskal-Wallis multiple comparison analysis. Error bars indicate s.d.; ****P < 0.0001; ns, not significant). **(H)** Touch-evoked assay reveals reduction in responsiveness upon mechanical stimulus in DMSO- and MG132-treated *fla* mutants, as shown in the motion trace diagram. **(I)** Quantification of the total swim distance shows significantly reduced motility in DMSO- and MG132-treated *fla* mutant embryos compared to controls (N = 3, n =3, P < 0.0001; One-way ANOVA followed by Kruskal-Wallis multiple comparison analysis. Error bars indicate s.d.; **** P < 0.0001. ns: not significant). **(J-L)** Myh protein levels in MG132-treated *fla* embryos are restored and comparable to the wild-type situation (N=3, mean ± S.D, P =0.3739 determined using two-tailed t-test. Error bars indicate s.d.; ns P>0.05) **(J, K)**, whereas the levels of methylated K35 Myh were still significantly reduced in MG132-treated *fla* mutants (N=3, mean ± S.D, P=0.0004 determined using two-tailed t-test. Error bars indicate s.d.; *** P < 0.001) **(J, L)**.

Subsequently, to assess whether interference with the UPS protein degradation machinery might lead to the restoration of Myh levels and thereby to the rescue of thick filament assembly in Smyd1-deficient *fla* mutant embryos, we treated *fla* mutants with MG132 and assessed heart and skeletal muscle structure and function by birefringence analysis and *in vivo* video microscopy. Highly organized muscle tissue such as skeletal trunk muscle in zebrafish is able to polarize light resulting in a distinct birefringence signal. As shown in Fig. 3E, we found the disorganization of the skeletal muscle tissue in *fla* mutant zebrafish embryos as visualized by the severe reduction of the birefringence signal compared to a strong birefringence signal in wild-type siblings (Fig. 3E). Interestingly, treatment of *fla* mutant embryos with the UPS inhibitor MG132 does neither improve fractional shortening (N = 3, n = 4/5, p < 0.0001. sib: 35.31±3.75%; sib MG132: 35.82±5.37%; mut: 0.0±0.0%; mut MG132: 0.0±0.0% at 72 hpf) (Fig. 3F) nor myofibrillar organization as shown by birefringence analyses (N = 3, n = 4/5, p = 0.0001 (Fig. 3G). Additionally, we assessed the touch-evoked escape response (TEER) of *fla* mutant embryos treated with MG132 or DMSO and found that both, Smyd1-deficient *fla* embryos treated with MG132 or the DMSO-treated control showed significantly impaired motility quantified by the total swim distance (N = 3, n = 3, p < 0.0001. sib: 2.61±1.55 cm; sib MG132: 1.64±0.8 cm; mut: 0.04±0.03 cm; mut MG132: 0.06±0.04 cm at 72 hpf) (Fig. 3H, I). Finally, we assessed Myh levels in *fla* mutant embryos after MG132 treatment and found that after inhibiting proteasomal degradation by MG132, the amount of muscle Myh in *fla* mutants was significantly increased and comparable to Myh levels in wild-type siblings (N=3, P=0.3739) (Fig. 3J, K). However, K35 methylated Myh was not detected in MG132-treated Smyd1-deficient *fla* zebrafish embryos (N=3, P=0.0004) (Fig. 3J, L).

These findings suggest that the Smyd1-mediated methylation of Myh at K35 is fundamental for the orchestrated assembly of the thick filament Myh into functional sarcomeric units.

### SMYD1-Mediated Methylation of Myosin K35 is Conserved in Human Cardiomyocytes

To investigate whether Smyd1-mediated Myosin K35 methylation is also critical for thick filament assembly in human striated muscle tissue, we next ablated SMYD1 by using Adeno-associated virus (AAV-) mediated SMYD1-specific shRNA in human iPSC-derived ventricular cardiomyocytes (hiPSCMs) and assessed total MYH levels, MYH K35 methylation as well as myofibrillar organization.

AAV1-mediated SMYD1-specific shRNA delivery (AAV1-SMYD1) into hiPSCMs resulted in the efficient knock-down of endogenous SMYD1 (N=4, P=0.0007) (Fig. 4A, B) and subsequently the significant reduction of MYH levels (N=5, P=0.0079) (Fig. 4A, C) compared to the Scrambled control AAV1 shRNA and similar to the observed situation in Smyd1-deficient zebrafish embryos. We next assessed MYH K35 methylation in AAV1-SMYD1 treated hiPSCMs and found the significant reduction of K35 methylation after SMYD1 knock-down (N=4, P=0.0051) (Fig. 4D, E), substantiating that this biological phenomenon is conserved between zebrafish and humans and further implying a fundamental role during thick filament assembly.

**Figure 4.**
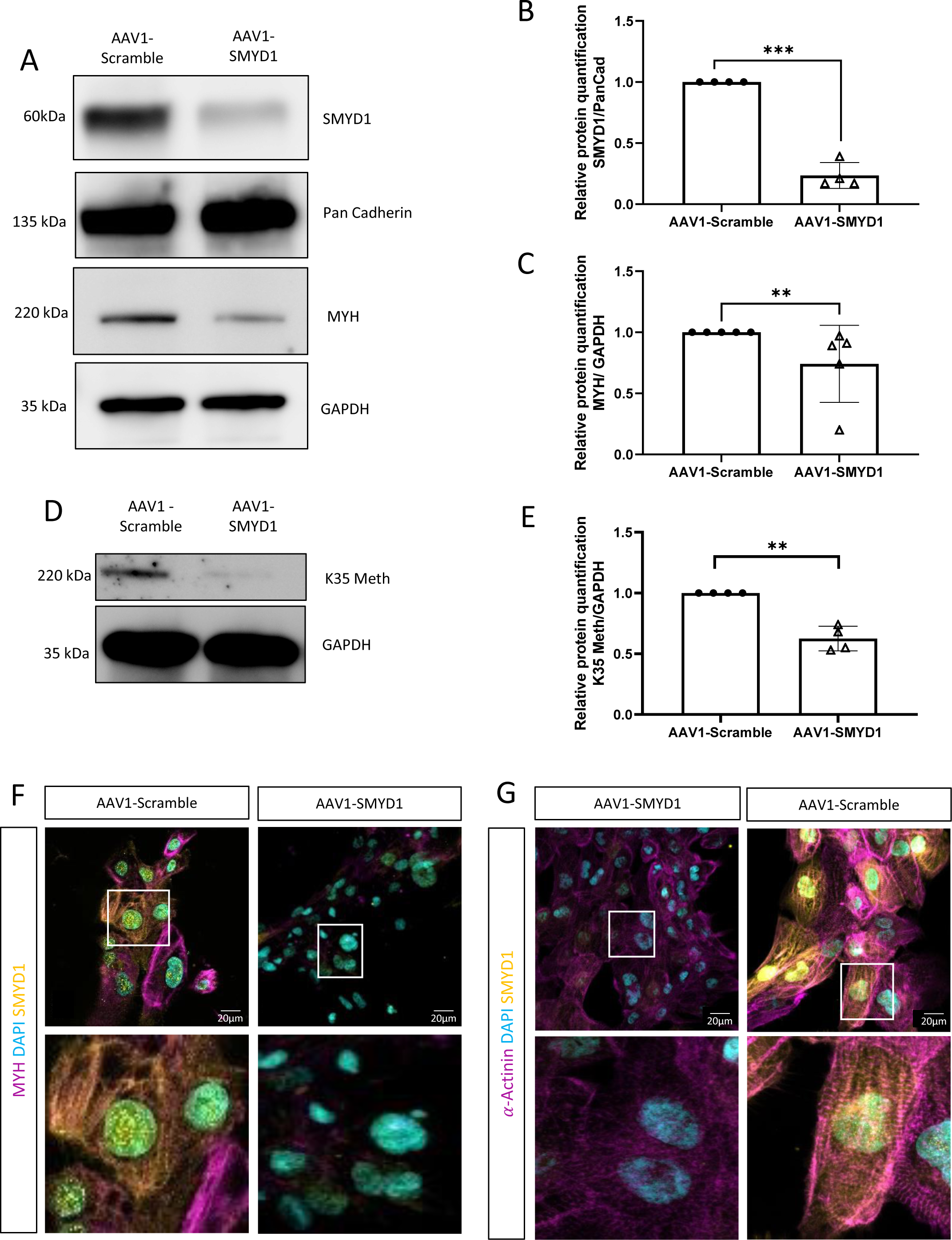
SMYD1-mediated methylation of MYH K35 is conserved in human iPSC-derived cardiomyocytes. **(A-C)** AAV1-SMYD1-mediated ablation of SMYD1 iPSC-derived cardiomyocytes leads to the significant downregulation of Myosin protein levels compared to control (AAV1-Scramble) treated cardiomyocytes. Pan Cadherin and GAPDH were used as loading control, respectively. (SMYD1: N=4, mean ± S.D, P =0.0007 determined using two-tailed t-test. Error bars indicate s.d.; *** p < 0.001) (MYH: N=5, mean ± S.D, P =0.0079 determined using two-tailed t-test. Error bars indicate s.d.; ** P < 0.01). **(D-E)** SMYD1-mediated Lysine 35 (K35) methylation is significantly reduced in AAV1-SMYD1-treated human iPSC-derived cardiomyocytes (N=4, mean ± S.D, P =0.0051 determined using two-tailed t-test. Error bars indicate s.d.; ** P < 0.01). **(F)** Immunostaining performed on 45 days old AAV1-SMYD1-treated cardiomyocytes demonstrated the absence of MYH (magenta) and SMYD1 (yellow) accompanied by the disruption of sarcomeric organization. Cell nuclei are shown by DAPI staining (cyan). **(G)** AAV1-SMYD1-treated cardiomyocytes show the severe reduction of the levels of K35 Myosin methylation (yellow) and the disorganization of sarcomeric structures as demonstrated by α-Actinin staining. Cell nuclei are depicted by DAPI staining (cyan).

Finally, we analysed whether the knock-down of SMYD1 and the subsequent loss of MYH K35 methylation leads to defective thick filament assembly and myofibrillar disarray, we performed immunofluorescent stainings for sarcomeric α-Actinin. Similar to the situation in zebrafish, we found pronounced myofibrillar disorganization in human iPSC-derived cardiomyocytes after SMYD1 deficiency mediated MYH K35 methylation loss (Fig. 4F, G).

In summary, our findings demonstrate that Myosin methylation at lysine 35 is specifically processed by SMYD1 and that MYH K35 methylation is critical for thick filament assembly in zebrafish and humans.

## Discussion

Methylation of lysine residues on histones but also non-histone proteins is a widespread post-translational protein modification that controls a wide range of biological phenomena. In this context, histone methylation in cell nuclei is a well-known and critical modification to control and fine-tune gene expression, DNA replication and also DNA repair [14]. However, the biological impact of non-histone protein methylation, particularly in the cytoplasm, as well as the specific methyltransferases that mediate this modification are not sufficiently understood. Here, we demonstrate that Smyd1, a methyltransferase that is expressed in both, the nucleus and the cytoplasm of striated muscle cells, mono-methylates sarcomeric Myh at lysine 35 thereby critically regulating thick filament assembly and sarcomerogenesis in zebrafish and human iPSC-derived cardiomyocytes.

The five members of the SET and MYND domain containing protein family (SMYD1-5) are well-known histone methyltransferases (HMTs) that modulate gene transcription by methylating unacetylated lysine residues on histone tails [15–17]. Furthermore, different members of the SMYD family are also involved in non-histone protein methylation thereby regulating their activity, function or stability in diverse cell types including skeletal muscle cells and cardiomyocytes[18, 19]. For instance, Smyd2 was shown to methylate the molecular chaperone Hsp90 in the cytoplasm of striated muscle cells which is important for the formation of a complex with the giant sarcomeric protein Titin [3]. Remarkably, Smyd2 ablation resulted in the loss of Hsp90 methylation, Titin instability and subsequently significantly impaired function of striated muscle [6]. Smyd2 was found to specifically mono-methylate Hsp90 at lysine residue K616 leading to their colocalization to the sarcomeric I-band. The striated muscle-specific methyltransferase Smyd1 was recently described to catalyse the methylation of histone 3 lysine 4 (H3K4) [20] as well as the site-specific lysine methylation of the heart and skeletal muscle-specific variant of nascent polypeptide-associated complex (skNAC) (K1975) [21]. skNAC K1975 methylation was found to control the activation of myoglobin (Mb) transcription, a striated muscle-specific hemoprotein important for oxygen transportation [21]. Interestingly, by using immunoprecipitation experiments and a Mass spectrometry-based approach we here identified muscle Myosin Heavy Chain as physical Smyd1 interactor and as another direct methylation target of Smyd1. We found that Smyd1 interacts with the head domain of Myh (1-111aa) which is highly conserved across isoforms and species and which is responsible for the binding of the Myosin ATPase and Actin. Although Smyd1-Myh interaction and the Smyd1-mediated methylation of Myosin was shown before [9], neither the Myh interaction domain nor the specific site of Myh methylation was known so far. In 2013, Li et al. were able to show that Smyd1b is able to methylate Myosin and that this could possibly have an influence on the stability of the protein [9]. Here, we describe the Smyd1-mediated mono-methylation of Myh at lysine 35 (K35) *in vivo* and *in vitro*. Although the methylation of this lysine residue has been the subject of speculation for over 40 years [22, 23], this post-translational modification has not yet been proven biochemically. The biological function of this methylation was also completely unclear. We were able to show that the failure to mono-methylate Myh K35 leads to the degradation of Myh and thereby induces severe defects in sarcomerogenesis in cardiomyocytes and skeletal muscle cells. Normally, the thick filament Myosin is incorporated into the sarcomeres of striated muscles with the help of the Myosin chaperones Hsp90a and Unc45b, thereby ensuring unhindered muscle development and function [7, 12]. We have now found that only correctly methylated Myh is incorporated into sarcomeres and unmethylated Myh is effectively degraded by the ubiquitin-proteasome system (UPS). Interestingly, a wide variety of Myosin mutations including haploinsufficiency inducing mutations are described to cause hypertrophic cardiomyopathy (HCM) in human patients. About 40 percent of HCM-causing mutations are found in the human cardiac Myosin gene and 1 in 500 people is suffering by HCM [24]. Although numerous HCM-inducing mutations in the N-terminal region of myosin have been described, no mutations of the Myh methylation site K35 have been described so far, so it is unclear whether altered Myh methylation contributes to the development of the disease [25]. Nevertheless, it is noteworthy that mutations in the human SMYD1 gene are also associated with the development of HCM [26, 27]. We have shown here that Smyd1 mono-methylates Myh at lysine K35 and that the loss of Smyd1 in zebrafish striated muscle but also in human iPSC-derived cardiomyocytes leads to severe defects in thick filament assembly, resulting in severely impaired muscle function. As shown by us and others, Smyd1 is exclusively expressed in cardiomyocytes and skeletal muscle cells in mice and zebrafish [7, 9, 28]. Targeted knockout of Smyd1 in mice results in lethality during early embryogenesis (at E10.5), due to impaired development of the right ventricular chamber and left ventricular contractile failure [26, 28]. In zebrafish, we found that SMYD1 is fundamental for thick filament assembly exclusively in cardiomyocytes and fast-twitch skeletal muscle cells und that the complete loss of Smyd1 function results in defective thick filament assembly due to the degradation of Myosin. It has not yet been investigated and is therefore unclear whether heterozygous mutations of Smyd1 in cardiomyopathy patients leads to pathologically altered Myosin methylation as well as stabilization and therefore leads or contributes to the onset or development of HCM.

HCM-causing variants of Myosin frequently alter the relaxed state of sarcomeric proteins resulting in elevated cardiomyocyte contractility, energy consumption and impaired left ventricular relaxation and filling [25]. Mavacamten, a selective small molecular cardiac Myosin inhibitor, which represents the first disease-specific treatment for HCM targeting the pathomechanism of the disease was approved by the FDA for adults with symptomatic HCM [29, 30]. Experimental and preclinical studies have shown that Mavacamten reduces the likelihood of Myosin-Actin cross-bridge formation by reducing the number of Myosin heads that can enter the “on-Actin” state and shifts the Myosin population to an energy-sparing, super-relaxed “off-Actin” state through binding the Myosin ATPase. Additionally, structural studies have shown that HCM mutations disrupt normal interactions between sarcomeric proteins, while Mavacamten normalizes these interactions, thereby reducing sarcomeric hyperactivity and hypercontractility. More studies are fundamental to assess whether Mavacamten is able to modify the pathomechanisms induced by other HCM-causing gene variants such as SMYD1 mutations that alter the methylation state of Myosin.

Non-histone methyltransferases, such as the Smyd family, are increasingly recognized as critical regulators of important biological processes, also in striated muscle cells. In this context, Smyd proteins such as Smyd1 and Smyd2 were recently shown to be fundamental for the proper assembly and stability of the sarcomeres, the basic contractile units of all striated muscle cells. We now show for the first time that Symd1-mediated K35 mono-methylation of sarcomeric Myosin heavy chain (Myh) is critical for regular thick filament assembly and that loss of K35 methylation leads to the proteasomal degradation of Myh and thereby severe sarcomeric defects in zebrafish but also human iPSC-derived cardiomyocytes. Future studies are now needed to investigate the pathological effects of defective Myh methylation, in particular on the development of HCM, and then to develop therapeutic strategies to offer new treatment options to patients with HCM.

## Materials and Methods

### Zebrafish strains

The present study was performed after appropriate institutional approvals (Tierforschungszentrum (TFZ) Ulm University, No. z183), which conform to EU Directive 2010/63/EU. Care and breeding of zebrafish, Danio rerio, was conducted as described previously [31].

### MG132 treatment

24 hpf *fla* mutant and wild-type sibling embryos were treated with 100 µM MG132 (Tocris Bioscience, Bristol, BS11 9QD, UK) in E3 for 24 h [32]. Similar amounts of DMSO were used as control.

### Zebrafish protein lysate and Western Blot Analysis

Western Blot analysis, protein lysates were prepared from 72 hpf from wild-type siblings and *fla* mutant embryos. 30µg of protein lysate were boiled in 5xLaemmli Buffer and separated on a SDS-PAGE (8-16%) and blotted onto polyvinylidene difluoride (PVDF) membranes. Then, the membranes were blocked using 5% milk powder in TBST for 1h at RT and incubated with primary antibodies overnight at 4°C. The membranes were washed and incubated with polyclonal anti-rabbit-IgG or anti-mouse-IgG antibody conjugated to horseradish peroxidase. All membranes were developed using the Pierce ECL Western Blotting Substrate (Thermo Scientific) and a luminescent image analyzer (Image Quant Las4000 mini) [33].

Antibodies used: mono-Methylated Lys35 antibody (Biogenes custom made, designed to detect the mono-methylated Lysine 35 of Myosin heavy Chain) (1:500 in 5% BSA), beta-actin (Cell Signalling 12620) (1:1000 in 5% milk), EB165 (Myosin Heavy Chain, embryonic and adult fast) (Hybridoma Bank) (1:500 in 5% milk), Ubiquitin (Cell Signalling 3936S) (1:1000 in 5% milk)

### Zebrafish immunostaining

72 hpf wild-type and *fla* mutant embryos were euthanized with tricaine, fixed in 4% paraformaldehyde (PFA) overnight at 4 °C, and embedded in 4% low melting agarose (melting temperature ≤65°C; Sigma) dissolved in distilled water. Longitudinal sections were cut with a Leica VT1200S vibratome to a thickness of 100μm and incubated in CAS-block solution (008120, Thermo Fisher) for 30 minutes (diluted 1:10 in distilled water). Staining was conducted in CAS-block (diluted 1:10 in distilled water). Images were acquired using Leica DMi8 confocal microscopes (Leica Mikrosysteme Vertrieb GmbH, Wetzlar, Germany).

Antibody used: EB165 (Myosin Heavy Chain, embryonic and adult fast) (Hybridoma Bank) (1:50), mono-Methylated K35 overnight (1:100) at 4°C followed by 1:100 Alexa-Fluor 488 and Alexa-Fluor 555 (both Invitrogen) for 2 h at RT.

### Birefringence analysis and heart and skeletal muscle functional assessment

Images were taken with a Zeiss Axio Zoom V.16 microscope and movies were recorded with a Leica DM IL LED microscope. Cardiac contractility was analyzed by assessing ventricular fractional shortening at 72 hpf as described previously [34]. Birefringence analysis was performed as previously described [34, 35].

### Touch-Evoked Escape Response Assay (TEER)

A TEER assay was performed as previously described [35] to analyze the motility of zebrafish embryos. E3 medium was maintained at 28 °C in a 14 cm diameter Petri dish (VWR), into which 72 hpf embryos were placed in the centre of the dish individually. The embryos were exposed to daylight for 10 min prior to this to allow them to acclimate. After placement, embryos were touched with a needle at the tip of the caudal fin. From the point of stimulus, a 10 s video was recorded using a digital camera (1080 pixels and 60 frames per second). Semi-automated motion tracking analysis of the obtained videos was performed with Tracker Video Analysis and Modeling Tool open-source software (physlets.org, accessed on 7 February 2023). The head of each embryo was used as the centre of mass. Tracking analysis was used to generate motion trace diagrams, and total swim distance and maximum velocity and acceleration were extracted from motion tracking data [35].

### Mass Spectrometry

Samples were prepared for mass spectrometric analysis following tryptic digestion. Employing an LTQ Orbitrap Velos Pro system (Thermo Fisher Scientific, Bremen, Germany) online coupled to an U3000 RSLCnano (Thermo Fisher Scientific, Idstein, Germany), samples were analyzed as described previously [36]. Database search was performed using MaxQuant Ver. 1.6.3.4 (www.maxquant.org) [37] by correlating fragmentation spectra with the UniProt zebra fish reference proteome set (www.uniprot.org) for peptide identification. Carbamidomethylated cysteine was considered as a fixed modification. In addition to methylated lysine, oxidation (M), and acetylated protein N-termini were allowed as variable modifications. False Discovery rates were set on peptide, modification and protein level to 0.01.

### Expression and purification of recombinant SMYD1

SMYD1 was expressed as GST-HIS-tagged recombinant protein. The production and purification were outsourced to Trenzyme (Trenzyme GmbH).

### pV5-DEST Smyd1b and pDEST53 Myh purification and overexpression

The Smyd1b and N-terminal Myh (1-111aa) coding sequence were respectively cloned into the pV5-DEST and pDest53 (expressing a GFP tag) vectors and transformed into One Shot TOP10 bacteria (Thermo Scientific, Waltham, MA). After overnight growth, plasmid purification was done using the QIAprep Spin Miniprep Kit (QIAGEN GmbH).

### Recombinant N-term MYH

N-term MYH1 (1-111 aa) peptide was synthetized by BioCat and customized with different tags: His-; Flag-. BioCat synthetized also the Flag-N-term MYH peptide with lysine replaced by alanine at position 35 (K35A).

To note: Lysine at position 35 (K35) in the human and zebrafish skeletal muscle Myosin heavy chain correspond to lysine at position 34 (K34) in the human heart-specific Myosin heavy chain 7 (MYH7).

### Immunoprecipitation assay

HEK293T cells were transfected with 1µg of pV5-DEST Smyd1b and pDEST53 (expressing GFP tag) Myh (1-111 aa) plasmid DNA using polyethylenimine. After 48 hours, GFP-tagged Myh was isolated from cell lysate using Dynabeads Protein G (Thermo 10004D) and GFP-tag Antibody (Roche #11814460001). IP and input were boiled in 5xLaemmli Buffer and separated on an SDS-PAGE (8-16%) and blotted onto polyvinylidene difluoride (PVDF) membranes. Then, the membranes were blocked using 5% milk powder in TBST for 1h at RT and incubated with primary antibodies overnight at 4°C. The membranes were washed and incubated with polyclonal anti-rabbit-IgG or anti-mouse-IgG antibody conjugated to horseradish peroxidase. All membranes were developed using the Pierce ECL Western Blotting Substrate (Thermo Scientific) and a luminescent image analyzer (Image Quant Las4000 mini) [33].

Antibodies used: V5-tag (cellsignaling #13202) (1:1000 in 5% milk), Anti-GFP (abcam #Ab12181) (1:1000 in 5% milk).

### In vitro methyltransferase experiments

4µg GST-His-SMYD1 were incubated at 37°C with 4µg synthetic peptide substrates corresponding to the 111 N-terminal amino acids Myosin Heavy Chain peptide (supplied by BioCat) in presence of 40µM s-adenosyl methionine (SAM) in in MTase assay buffer (50 mM Tris–HCl [pH 8.5], 20 mM KCl, 10 mM MgCl2, 10 mM β-mercaptoethanol, and 1.25 M sucrose). Reactions were stopped after 1 hour boiling the sample at 70°C. Methylation signal was then detected with antibodies against Methylated Lysines (abcam/ab23366) and mono-Methylated K35 (custom made by Biogenes), together with His tag (cell signalling #2365) and Flag tag (cell signalling #8146) signal detection.

### IPSC-derived cardiomyocytes

IPSC-derived ventricular cardiomyocytes were differentiated as previously described [38]. Experiments were performed with 45 days old iPSC-derived ventricular cardiomyocytes.

### AAV1-shRNA SMYD1 and AAV1 Scramble infection of iPSC-derived cardiomyocytes

To knockdown SMYD1, the cardiomyocytes were infected using pAAV[shRNA]-EGFP-U6>Scramble_shRNA as control and pAAV[shRNA]-EGFP-U6>hSMYD1_shRNA (VectorBuilder) with a MOI of 50.000. Briefly, 36 days old cardiomyocytes were seeded in a multiwall plate, after they recover and start beating, usually 40 days old the cells were infected with AAV1 using the MOI of 50.000. After 24 h the virus-containing medium was aspirated and changed with fresh cardiomyocytes maintenance medium. Transfection expression was observed from 24 hours post-infection by green fluorescent signal using the Leica DMIL LED and remained detectable for more than a week in the non-dividing cardiomyocytes. 4 days after infection further experiments were performed (Western Blot, immunostaining).

### iPSC-derived ventricular cardiomyocytes protein extraction

The cardiomyocytes were harvested and collected in a 50 ml conical tube. Next, the suspension was centrifuged at 100 g for 10 min and the supernatant was removed. The pellet was resuspended in 5 ml of DPBS and then centrifuged at 100 *g* for 10 min. After aspirating the supernatant, the pellet was resuspended in RIPA buffer (supplemented with protease inhibitor). The suspension was transferred to 1.5 ml tubes and kept on ice for 15 min. Next, the cells were centrifuged at 4°C and 13200 rpm for 15 min and then the supernatant was collected in a new 1.5 ml tube. Finally, the concentration was determined by BCA-assay and the protein was stored at −80°C.

### iPSC-derived ventricular cardiomyocytes Western Blot

The protein lysates of the SMYD1 scramble RNA and shRNA (10 µg) were boiled with 5x Laemmli buffer at 95°C for 5 min. Next, the protein lysates were loaded into a precast 4-20% protein gel, they were separated by SDS-PAGE (for 1 Gel: constant 35 mA, 45 min), and blotted onto a polyvinylidene difluoride (PVDF) membrane (constant 100 V, 1 h). Then, the membrane was blocked in 5% BSA in TBST at room temperature and afterward incubated with the primary antibody at 4°C overnight. On the next day, after washing, the membrane was washed four times, for 10 min each, and incubated with the analogous secondary antibody (polyclonal HRP-linked IgG anti-rabbit or anti-mouse) for 1 h at room temperature. Afterwards, the membrane was washed again four times, for 10 min each. Signal development and quantification were administered by the usage of Pierce ECL Western Blotting Substrate (Thermo Fischer Scientific) and an ImageQuant LAS4000 mini.

Antibodies used: mono-Methylated Lys35 antibody (Biogenes custom made) (1:500 in 5% BSA), GAPDH (abcam, ab181602) (1:1000 in 5% milk), MF20 (MYH) (Hybridoma Bank) (1:500 in 5% milk), SMYD1 (Thermo, PA5-31482) (1:5000 in 5%milk)

### Immunostaining on iPSC-derived ventricular cardiomyocytes

For immunostaining, the cardiomyocytes were plated on glass bottom plates. 4 days post-AAV1 infection of the cells, they were washed in DPBS with 0.02% sodium acid (DPBS+) and then fixed in 4% PFA in DPBPS+ at room temperature for 20 min. After washing, the cells were incubated in 0.5% Triton X-100 at room temperature for 5 min and then washed in DPBS+ twice. For blocking, 1x power blocking buffer was used, and the fixed cardiomyocytes were blocked at room temperature for 30 min. Afterwards, the cells were incubated with the primary antibody in 1x blocking buffer at 4°C overnight. On the next day, the cardiomyocytes were washed twice in DPBS+ at room temperature for 5 min, followed by the incubation with the corresponding secondary antibody (Goat Anti-rabbit IgG Alexa Fluor 488, Goat Anti-mouse IgG1 Alexa Fluor 555, Goat Anti-mouse IgG2b Alexa Fluor 555) in 1x blocking buffer at room temperature for 1 h. After two more washing steps in DPBS+ for 5 min, DAPI in DPBS+ (1:5000) was added to the cells and they were then visualized by using a Leica TCS SP8.

Antibodies used: mono-Methylated Lys35 antibody (Biogenes custom made) (1:200), MF20 (Hybridoma Bank) (1:10), SMYD1 (Thermo, PA5-31482) (1:100).

### Statistical analysis

Data were analysed using GraphPad Prism 9 software. All experiments were conducted in minimum biological triplicates (N = 3). All the results are expressed as mean ± standard deviation (S.D.) and statistical analyses were conducted as described in the figure legends. A p-value smaller than 0.05 was taken as statistically significant.

## Acknowledgements

We would like to thank Regine Baur, Renate Durst, Karin Strele, Katrin Vogt, Sabrina Diebold, Jessica Hofmiller and Denise Mann for their excellent technical assistance.

We thank Dr. Sebastian Wiese and the Core Unit Mass Spectrometry and Proteomics (Ulm University) for the Mass-Spectrometry analysis.

This work was supported by the Deutsche Forschungsgemeinschaft (DFG) JU2859/7-1 and JU2859/9-1 (SJ) and the German Federal Ministry of Education and Research (BMBF) (e:Med-SYMBOL-HF grant #01ZX1407A; e:Med-coNfirm grant #01ZX1708C) (SJ). The funders had no role in study design, data collection and analysis, decision to publish, or preparation of the manuscript.

---

**Figure S.1.**
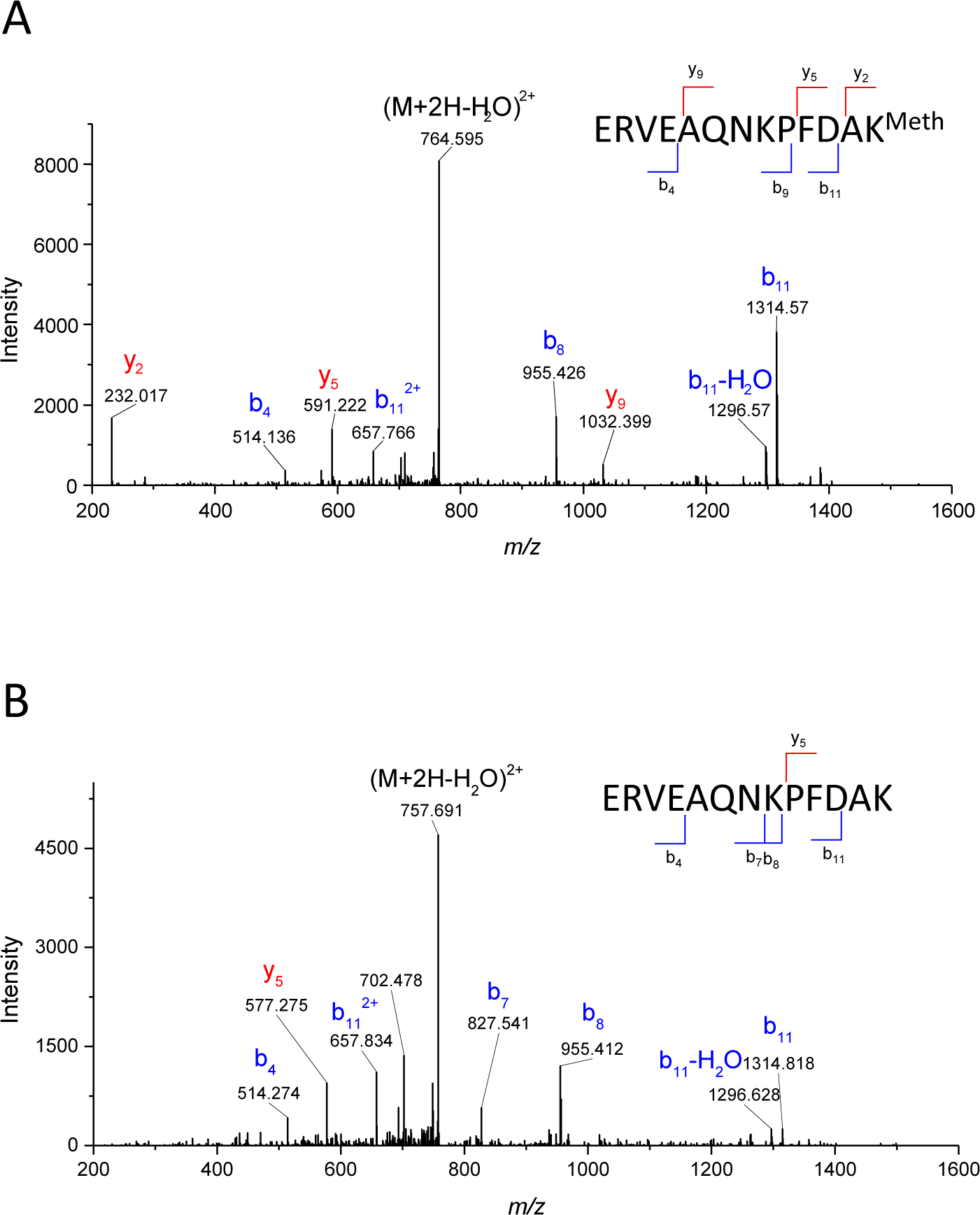
MS fragmentation spectra of K35-methylated (A) and non-methylated (B) MYH peptide amino acids 23 to 35. It is possible to observe b8-b11-, y2- and y5-ions in the respective MS2 spectrum, mapping the methylation to K35.

**Figure S.2.**
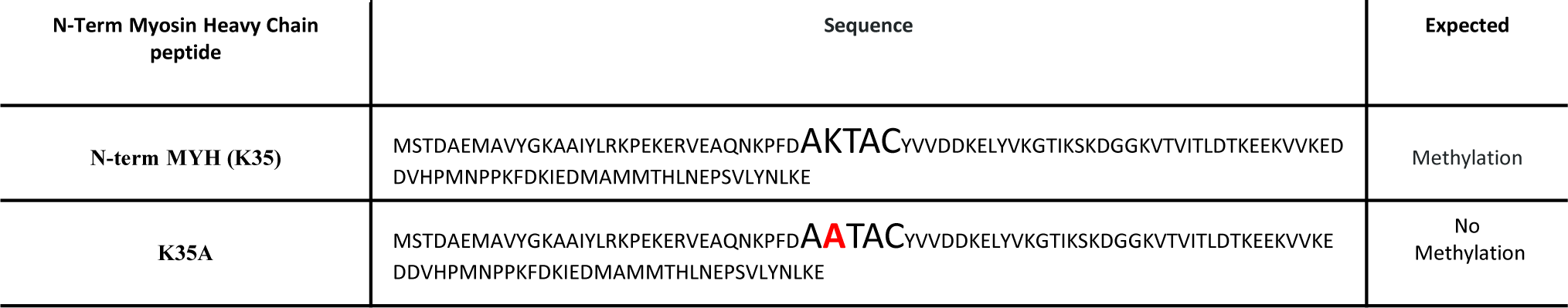
Sequences of the N-term MYH and the mutated sequence on N-term MYH with the Lysine to Alanine exchange at position 35.

